# A mouse model of insomnia with sleep apnea

**DOI:** 10.1101/2022.08.16.503987

**Authors:** Satoru Masubuchi, Takako Yano, Kouji Komatsu, Keisuke Ikegami, Wataru Nakamura, Akinobu Ota, Sivasundaram Karnan, Kosei Takeuchi, Yoshitaka Hosokawa, Takeshi Todo, Toshiaki Shiomi

## Abstract

Obstructive sleep apnea (OSA) patients are exposed to nighttime hypoxia during sleep by intermittent airway closure and feel daytime strong sleepiness. Strangely, insomnia co-occur in some OSA patients, which is called co-morbid insomnia and sleep apnea (COMISA). Here, we show activity responses to daytime hypoxia (DHx) in nocturnal mice were comparable to daytime sleepiness and co-occurring nighttime insomnia in COMISA. DHx reduced activity in active phase (AP) and increased following activity in activity ending phase (AEP). This down-and-up activity response (DUR) by DHx was also observed in molecular clock deficient *Cry1* and *Cry2* double knockout mice (CryDKO) expressing nighttime activity rise under light-dark cycle (LD) and not observed in arrhythmic CryDKO under constant darkness (DD). When daytime timing hypoxia was exposed at transition from LD to DD, about every 6 h down and up and down wavelike activity responses appeared in arrhythmic CryDKO. Results indicate this wavelike response and AP activity overlap and cause DUR in rhythmic mice. DHx increased plasma corticosterone and this increase antagonized AP activity reduction by DHx. DHx reduced forebrain adenosine and morning adenosine inhibition by caffeine induced DUR. Adenosine inhibition by caffeine or istradefylline at transition from LD to DD induced wavelike response in CryDKO. It is possible that wavelike response is damped oscillation because, interestingly, chronic caffeine treatment induced circasemidian and/or circadian activity rhythms in arrhythmic CryDKO. Evening caffeine attenuated DUR by DHx, which suggested adenosine inhibition chronotherapy may improve OSA/COMISA symptoms. Our animal model will be useful to understand COMISA.

**Significance:** Obstructive sleep apnea patients (OSA) are exposed to nighttime hypoxia during sleep. OSA feels daytime strong sleepiness and increases risk of many diseases. Insomnia occurs in not a few OSA, which is called comorbid insomnia and sleep apnea (COMISA). We show here a mouse model of COMISA. In mice, daytime hypoxia exposure induced following down and up activity response (DUR), activity reduction in active phase and increase in activity ending phase, which corresponded to sleepiness and insomnia in COMISA. We found DUR was clock gene independent and might be driven by circasemidian system. Glucocorticoid and forebrain adenosine response were involved in DUR. Caffeine chronotherapy was effective in DUR. Our model may be useful to understand COMISA.

## Introduction

Circadian physiological and behavioral rhythms in mammals are driven by the molecular clock in the hypothalamic suprachiasmatic nucleus (SCN). The molecular clock system has a transcriptional-translational inter-relationship among clock genes [1]. Clock gene deficient mice, for example *Cry1* and *Cry2* double knockout mice (CryDKO) [2,3], show arrhythmic behavior under constant darkness (DD) [4] but they show nighttime activity increase in response to light dark cycle. Circadian time dependently, hypoxia affects mammalian physiology [5–7]. Nighttime hypoxia has a strong impact in humans. Nighttime hypoxia more reduced arterial oxygen saturation [8]. Symptoms of Acute Mountain Sickness [9] often develop or worsen after the first night of sleep at high altitude [10].

Obstructive sleep apnea (OSA) is a common sleep disorder. OSA patients are exposed to nighttime hypoxia during sleep by intermittent airway closure and feel daytime strong sleepiness. OSA increases risk of hypertension, stroke, heart failure, diabetes, car accidents, and depression [11]. Strangely, the comorbidity between insomnia and OSA is highly prevalent [12–16]. This is called comorbid insomnia and sleep apnea (COMISA) [13, 14, 16]. COMISA is difficult to treat only by standard treatment of OSA, continuous positive airway pressure. Advanced positive airway pressure mode, adaptive servo-ventilation [17, 18], and cognitive and behavioral therapy for insomnia [16, 19, 20] are applied for treatment of COMISA. COMISA is also called, sleep apnea plus [21], sleep-disordered breathing plus [22], complex insomnia [12, 23], OSA-insomnia [12, 24]. After the first report of insomnia with sleep apnea in 1973 [25], research reports about COMISA are increasing [14]. However, what drives this strange and paradoxical response, coexistence of sleepiness and insomnia, is still elusive.

We found hypoxia exposure at daytime (rest phase) reduced activity in active phase (AP) and increased following activity in activity ending phase (AEP) in mice. This down-and-up activity response (DUR) was comparable to daytime sleepiness and co-occurring nighttime insomnia in COMISA. Like nighttime apnea (hypoxia) in diurnal humans, daytime hypoxia (DHx) in nocturnal mice similarly affected active and rest phase behaviors. Therefore, we considered DUR by DHx is a mouse model to understand the pathophysiology of insomnia with sleep apnea. In this study, we analyzed DHx exposed mice to show the factors which regulate behavioral response to DHx.

## Materials and methods

### Animals

Specific pathogen free (SPF) male C57BL/6J and ICR mice were purchased from Japan SLC. *Cry1*−/− and *Cry2*−/− mice of C57BL/6J were generated from *Cry1*+/− and *Cry2*+/− mice [3] by backcrossing of ten generations with the mice of C57BL/6J strain [26]. Mice were maintained at the animal facilities of Aichi Medical University. The experimental protocols of this study were approved by the Committee for Animal Research at Aichi Medical University.

### Behavioral rhythm monitoring

Mice at 2-12 months of age were housed individually and their behaviors were monitored under 12 h light (fluorescent light, ~100 lux) - 12 h dark cycle (LD). The light off time was defined as Zeitgeber Time (ZT) 12. The locomotor activities of mice were monitored by passive (pyroelectric) infrared sensors (PS-3241; EK-Japan, Japan) [27]. Data were collected and analyzed as described previously [28] using The Chronobiology kit (Stanford Software Systems, Stanford, CA, USA). Mice were entrained to LD for more than two weeks and used for experiments. We calculated activity values for each mouse by taking the values of the amount of 24 h activity before hypoxia exposure as 1. We used these relative activity values for analysis. Circasemidian/circadian periods of activities were identified by chi-square periodogram [29].

### Hypoxia exposure and drug administration

For hypoxia (8% oxygen) exposure, the mouse cages in room air (21% oxygen) were moved from under the infrared sensors into air sealed transparent bags. Pure nitrogen was infused in the bag with monitoring of oxygen concentration by oxygen content meter (Oxy-1; JIKCO, Tokyo, Japan). After hypoxia exposure, animals were returned to under the infrared sensors.

For drug treatments by intraperitoneal injections, caffeine (15, 30 mg/kg), metyrapone (50 mg/kg) and istradefylline (3.3 mg/kg) were dissolved in vehicles as follows: Caffeine (FUJIFILM Wako 031-06792) was dissolved in saline. Metyrapone (CAYMAN CHEMICAL 14994) was dissolved in saline containing 20% EtOH. Istradefylline (Tokyo Chemical Industry I1100) was dissolved in saline containing 33% Dimethyl Sulfoxide (FUJIFILM Wako 046-21981).

For chronic caffeine treatment, caffeine (0.5g/L) was dissolved in distilled water and administrated as drinking water.

### Quantitative PCR analysis

Extraction of forebrain mRNA was performed as follows. Mice were sacrificed by cervical dislocation and forebrains were removed immediately, frozen on dry ice and stored at −80 □ until use.

Each forebrains were homogenized using the POLYTRON homogenizer (Kinematica AG, Switzerland) in 20 ml TRIzol RNA isolation reagents (Thermo Fisher Scientific 10296010). Total RNA isolated from TRIzol was reverse-transcribed using a SuperScript III First-Strand Synthesis System for RT-PCR (Thermo Fisher Scientific 18080-051) according to the manufacturers’ instructions. Quantitative PCR analysis (qPCR) of individual cDNA was performed using the DyNAmo HS SYBR Green qPCR Kit (Thermo Fisher Scientific F410L) employing the QuantStudio^®^ 3 Real-Time PCR systems (ABI). qPCR primers are shown in Table S1.

### Microarray gene expression analysis

We performed a comprehensive gene expression analysis in the ICR mice forebrains exposed to DHx (n=2) and controls (n=2). We used a Mouse Gene Expression 4 × 44K Microarray chip (G4846A, Agilent Technologies), which can examine 23,215 genes as described [30]. For the analysis of the gene expression profile, total RNA isolated using the TRIzol reagent was purified using the RNeasy Mini Kit (QIAGEN 74106) according to the manufacturer’s instructions. The quality of the isolated RNA was ascertained using a NanoDrop 1000 Spectrophotometer (Thermo Fisher Scientific). The experimental procedure for cDNA microarray analysis was performed according to the manufacturer’s instructions (Agilent Technologies) and signal intensities were normalized as previously described [31]. Hypoxic Forebrain Microarray dataset reported in this paper has been deposited into NCBI’s Gene Expression Omnibus under accession number, GEO: GSE135231.

### High-performance liquid chromatography (HPLC)

Adenosine and inosine contents of the forebrains were examined by reversed-phase HPLC system as described [32]. We used the Nexera UHPLC (SHIMADZU, Kyoto, Japan) system which included Solvent Delivery Unit (LC-30AD), Autosampler (SIL-10AP) and UV-Vis Detectors (SPD-20A). Analyte separation was achieved by using L-column 2 ODS (5 μm, 4.6×150 mm, Chemicals Evaluation and Research Institute, Japan 722070). UV-Vis Detection was set at a wavelength of 254 nm.

Extraction of forebrain samples for HPLC was carried out as follows. Mice were sacrificed by cervical dislocation and forebrains were immediately removed, frozen on dry ice and stored at −80 □. Forebrains were homogenized using the POLYTRON homogenizer in 20x volume of PE buffer containing 50 mM phosphoric acid and 0.1 mM EDTA (v:w). Homogenized samples were centrifuged at 16000 g for 15 min. The supernatants were filtered through a 0.22 μm Cellulose Acetate filter tube (Corning^®^ Costar^®^ Spin-X^®^ centrifuge tube filters cellulose acetate membrane; SIGMA-ALDRICH CLS8160) at 16000 g for 5 min and the filtrates were directly used for HPLC. Adenosine (SIGMA-ALDRICH A9251) and inosine (SIGMA-ALDRICH I4125) were dissolved in PE buffer and used as standards.

A dual mobile phase gradient was used. Mobile phase A contained 0.52 mM sodium 1-pentanesulfonate (Tokyo chemical industry I0343) and 0.20 M KH2PO4 (FUJIFILM Wako 169-04245) at pH 3.5 using 85% phosphoric acid (SIGMA-ALDRICH 345245). Mobile phase B had the same final concentrations as that of the mobile phase A, except for the addition of 10% acetonitrile (v/v). The gradient composition of mobile phase B was 0% at 0 min, 0% at 1 min, 100% at 6 min, 100% at 11 min, 0% at 12 min and 0% at 20 min. The flow rate was 1.0 mL/min and the sample injection volume was 50 μl.

### Enzyme-linked immunosorbent assay (ELISA)

Plasma corticosterone levels were examined using Corticosterone ELISA Kit (CAYMAN CHEMICAL 501320) according to the manufacturer’s instructions.

### Statistics

The significance of differences between various groups were analyzed by Student’s t-test, two-way ANOVA or one-way ANOVA. Statistical analyses were performed with Statcel software, OMS Publishing Inc., Japan

## Results

### DHx reduced activity in AP and increased activity in AEP

We examined circadian timing dependent hypoxia effects on daily locomotor activity. 9h hypoxias (8%) were exposed to ICR mice under LD (Figure 1A-I). In four timings examined, DHx (ZT0-9) (Figure 1A) and dawn hypoxia (ZT18-3) (Figure 1D) reduced the 24 h activity amounts after exposure (Figure 1E) and changed following activity variations (two-way ANOVA) (Figure 1F, I). Interestingly, DHx reduced the following AP activity (ZT12-18, ZT20-0) and increased the AEP activity (ZT2-4) (Figure 1F). This DUR did not occur by dawn hypoxia although AP activity was reduced (ZT12-16, ZT18-20) (Figure 1I).

**Figure 1.**
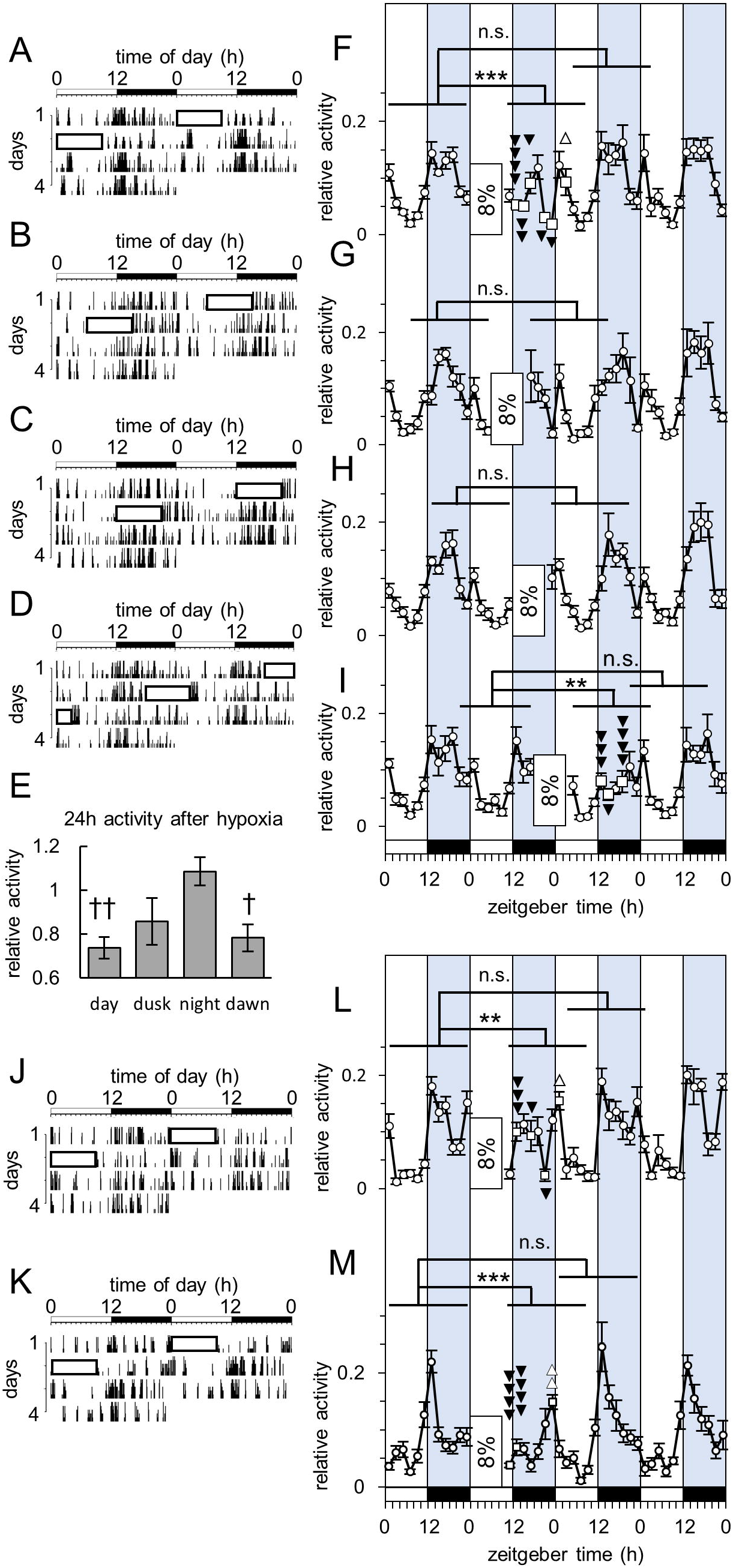
DHx under LD induced DUR not only in molecular clock-intact mice but also in CryDKO. (A-D, J, K) Representative double plotted actograms of ICR mice (A-D), WT (J) and CryDKO (K). The black and white bars above the recordings represent the LD cycles. Rectangles in the actograms show exposures to DHx (ZT0-9, 8%) (A, J, K), dusk hypoxia (ZT6-15, 8%) (B), nighttime hypoxia (ZT12-21, 8%) (C) and dawn hypoxia (ZT18-3, 8%) (D). (E) Relative amounts of activities from 1 h to 25 h after hypoxia corresponding to (A)-(D). We compared activity amounts of 24h before hypoxia and from 1 h to 25 h after hypoxia (DHx; n=8, dusk hypoxia; n=8, nighttime hypoxia; n=8, dawn hypoxia; n=8). †p<0.01, ††p<0.001, Student’s t-test. (F-I, L, M) Relative amounts of daily activities corresponding to (A)-(D), (J), (K): exposures to DHx (F; ICR, n=8, J; WT, n=8, K; CryDKO, n=10), dusk hypoxia (G; ICR, n=8), nighttime hypoxia (H; ICR, n=8) and dawn hypoxia (I; ICR, n=8). Open circles and open squares indicate relative values of every 2 h activities (mean±SEM). Open rectangles are hypoxia exposures. The black and white bars under the values represent the LD cycles. We compared these every 2h activity variations of 24 h before hypoxia and from 1 h to 25 h after hypoxia with matching ZTs (two-way ANOVA; **p<0.0005, ***p<0.0001). Open squares indicate significant value changes. ⍰⍰⍰⍰p<0.0001, ⍰⍰⍰p<0.0005, ⍰⍰p<0.005, ⍰p<0.05, ⍰⍰p<0.005, ⍰p<0.05, Fisher PLSD.

### Active phase activity in molecular clock-deficient mice under LD cycle respond to DHx in the same ways as molecular clock-intact mice

DUR was long time, more than 10 h, response and occurred only by DHx (Figure 1A-D, 1F-I), which provided an idea that biological clock was involved in DUR. DHx increased hypoxia responsive *Vegfa* mRNA in the forebrain (Supplementary Figure S1A). DHx also increased *Per1, Nfil3*, and *Npas2* mRNA and decreased *Dbp* and *Bmal1* mRNA (Supplementary Figure S1B), which suggested molecular clock [1] was involved in DUR. Therefore, we exposed hypoxia to CryDKO (C57BL/6J background) that shows activity increase in dark phase under LD and loses activity rhythm under DD [2,3]. In wild type C57BL/6J mice (WT), DHx reduced following AP (ZT12-14, ZT16-18, ZT20-22) and increased AEP (ZT0-2) activities (Figure 1J, L) as observed in ICR mice. In CryDKO, DHx suppressed following AP activity (ZT10-14) and induced a new activity bout which peaked at late nighttime (ZT22-0) (Figure 1K, M). Behaviors of all mice, i.e., ICR, WT and CryDKO responded in a similar manner to DHx under LD. DHx reduced AP activity and increased AEP activity. Persistence of DUR in CryDKO indicated the behavioral response to DHx was molecular clock system independent.

### Hypoxia of daytime hypoxia timing (HDHT) starting from transition from LD to DD induced wavelike behavioral response in arrhythmic CryDKO

Results under LD (Figure 1) raised a possibility that not molecular clock but daily activity variation was needed for DHx response. To test this, mice were moved to DD and exposed to hypoxia. WT activity variation did not change by moving them from LD to DD (two-way ANOVA) (Figure 2A, E). Hypoxia of daytime hypoxia timing (HDHT) from 24 h to 33 h after DD (9 h, 8%) suppressed activity corresponding to AP (ZT16-18, ZT22-0) and increased activity corresponding to AEP (ZT0-2) in DD similar with under LD (two-way ANOVA) (Figure 2A, E). Hypoxia is known to affect mammalian circadian rhythm phase [33]. Acceleration of re-entrainment to a new LD cycle by hypoxia is regulated by hypoxia inducible factor 1α [34]. But, effect of HDHT on circadian rhythm phase was not observed.

**Figure 2.**
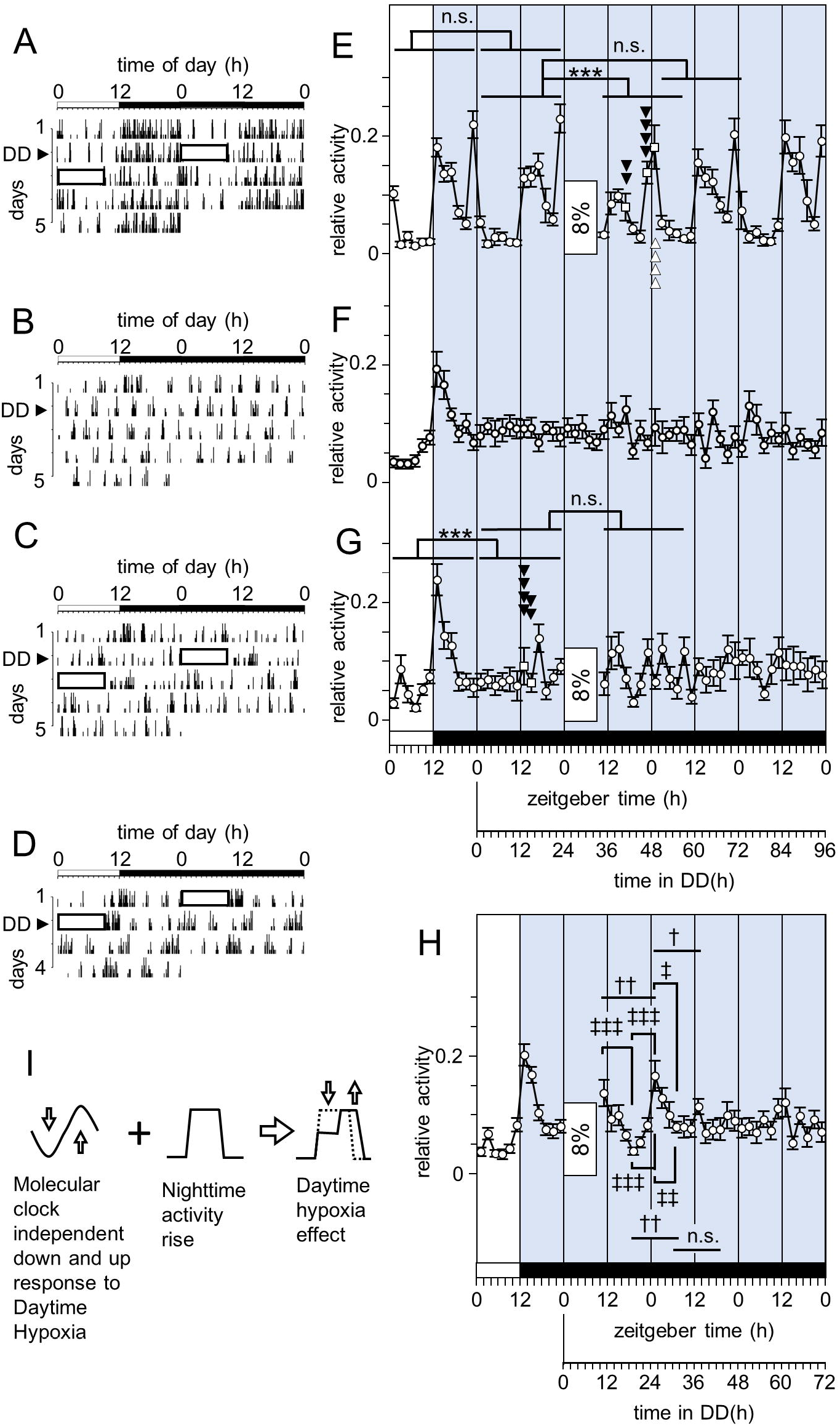
HDHT starting from LD-DD transition induced wavelike behavioral response in arrhythmic CryDKO. (A-D) Representative double plotted actograms of the transition from LD to DD of WT (A) and CryDKO (B-D). The black and white bars above the recordings represent the LD and DD. Black triangles indicate the day of moving from LD to DD. Rectangles in the actograms show HDHT (9 h, 8%) (A, C, D). Mice were exposed to HDHT after the first 24 h (A, C) in DD or from the beginning of DD (D). (E-H) Relative amount of daily behavioral activity corresponding to (A) to (D). Open circles and open squares indicate relative values of every 2 h activities (mean ± SEM). Open rectangles are HDHT (E, G, H). A group of CryDKO (n=14) (F) were moved from LD to DD and activities were monitored for 4 days without HDHT. The black and white bars under the values show the transition from LD cycles to DD. DD started from the end of dark phase of LD cycle. In WT (n=8) (E) and CryDKO (n=9) (G), 24 h activity variations before and after transition from LD to DD were compared (two-way ANOVA; ***p<0.0001). In same animals, and activity variations of 24 h before hypoxia (the first 24 h in DD) and from 1 h to 25 h after hypoxia with matching ZTs were compared (two-way ANOVA; ***p<0.0001). Open squares indicate significant value changes (Fisher PLSD; ⍰⍰⍰⍰p<0.0001, ⍰⍰p<0.005, ⍰⍰⍰⍰p<0. 0001). Activity down-and-up or up-and-down after HDHT in CryDKO (n=18) (H) were evaluated by one-way ANOVA (†† p<0.0005, † p<0.001). ‡‡‡ p<0.0001, ‡‡ p<0.001, ‡ p<0.01, Fisher PLSD. (I) Model of behavioral response to DHx.

Daily variation of activity in CryDKO under LD disappeared by moving from LD to DD (Figure 2B, F) as reported [2, 3]. After activity increase disappearance (ZT12-16) (two-way ANOVA), HDHT from 24 h to 33 h after DD did not affect following 24 h activity variation in CryDKO (two-way ANOVA) (Figure 2C, G). This result suggested that behavior responded to hypoxia when HDHT was exposed after activity bout. To test this, CryDKO were exposed to HDHT from the beginning of DD, the end of dark phase of final LD. After hypoxia, activity value decreased for 8 h from the first time point (10-12 h in DD) to 18-20 h time point, increased for next 6 h to 24-26 h time point, again decreased for next 6 h to 30-32 h time point and showed a small peak 6h later (36-38 h in DD) (Figure 2D, H). Variations in the first down and the first up (10-26 h in DD), in the first up and second down (18-32h in DD) and in the second down and second up (24-38h in DD) were significant (one-way ANOVA). But, variation in the second up and third down (30-44h in DD) were not significant (one-way ANOVA). Antecedent activity bout dependently, HDHT induced every about 6 h down and up and down response. Results indicated DHx induced clock gene independent short wavelike response. This wavelike response and nighttime activity bout overlapped and caused DUR in rhythmic mice (Figure 2I).

### Glucocorticoid secreted by DHx reduced the behavioral response to DHx

Next, we investigated forebrain at the end of DHx to know the factors which control the behavioral response to DHx. Microarray (Supplementary Figure S2A) and qPCR (Supplementary Figure S2B) analyses revealed that DHx increased a group of genes, *Cdkn1a, Fkbp5, SGK1, Tsc22d3,* and *Nfkbia,* expression levels, which reflected glucocorticoid (GC) action [35]. As expected from this, DHx increased plasma corticosterone robustly (Figure 3A). To know the role of the GC increase induced by hypoxia, metyrapone (MP; 50mg/kg) was administrated at the beginning of DHx (ZT0) (Figure 3B-E). MP reduced corticosterone level (ZT4) and suppression was over by the end of DHx (ZT9) (two-way ANOVA) (Figure 3F). The 24 h activity amount reduced markedly after hypoxia exposure with MP (two-way ANOVA) (Figure 3G). MP reduced activity of AP (ZT12-16, ZT22-0) in DHx exposed mice (two-way ANOVA) (Figure 3I). Interestingly, MP reduced activity of AP (ZT14-16) and increased activity of AEP (ZT2-4) (two-way ANOVA) without DHx (Figure 3H). These results indicated that endogenous GC increased by DHx attenuated the effect of DHx.

**Figure 3.**
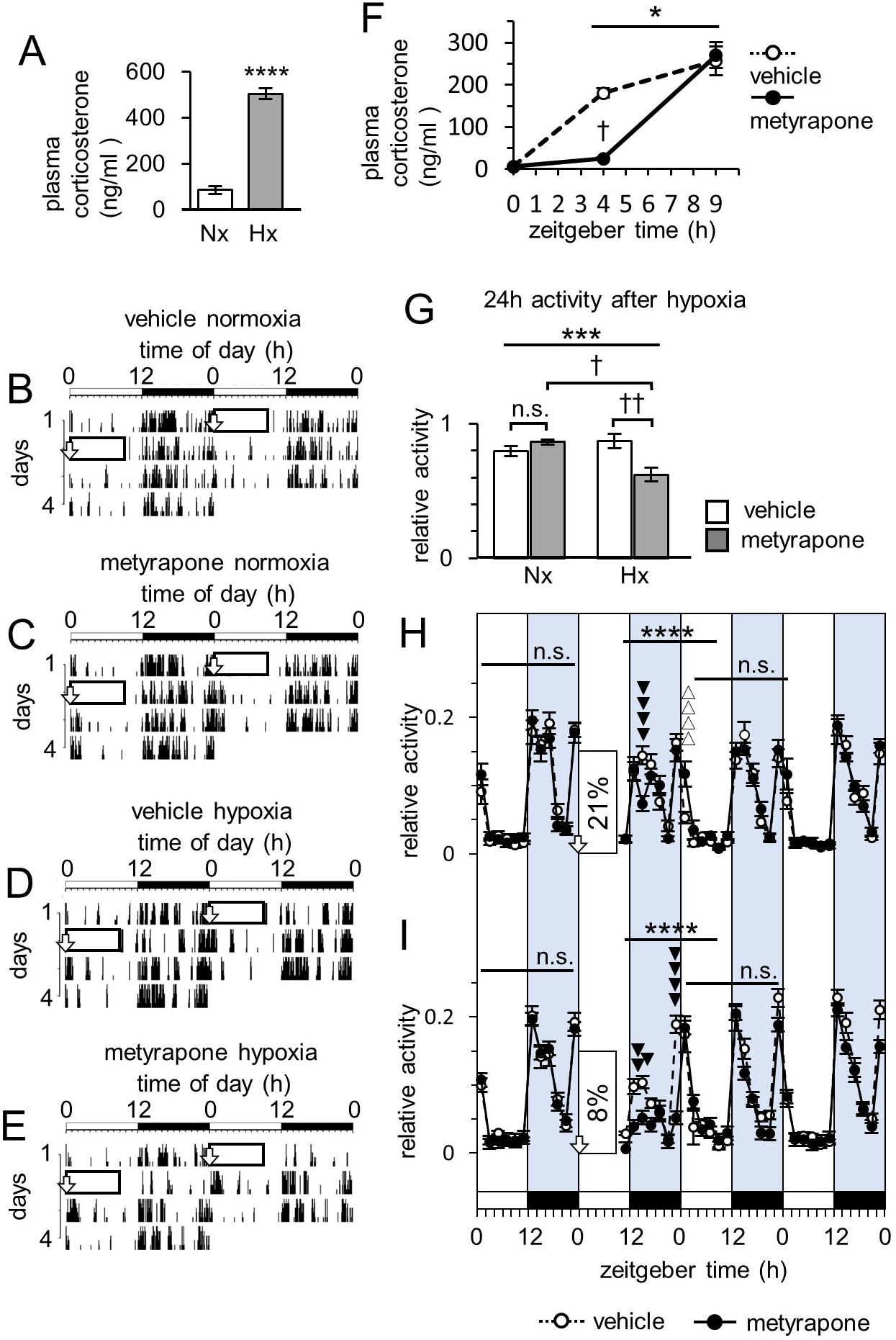
Involvement of GC on activity change by DHx. (A) Plasma corticosterone levels of ICR mice at the end of DHx (ZT9) (Hx; n=6) and normoxic control (Nx; n=6). ****p<0.0001, Student’s t-test. (B-E) Representative double plotted actograms of WT. The black and white bars above the recordings represent the LD cycles. Rectangles in the actograms show DHx (ZT0-9, 8%) (D, E) or normoxic control (21%) (B, C). Open arrows show vehicle (B, D) or MP (50 mg/kg) (C, E) at the beginning of DHx. (F) Plasma corticosterone levels of vehicle or MP treated (ZT0) and DHx-exposed WT at ZT0 (n=4), ZT4 (vehicle; n=5, MP; n=4) and ZT9 (vehicle; n=6, MP; n=6). We evaluated interaction between the timing effect (ZT4 or ZT9) and MP effect (two-way ANOVA; *p<0.01). †p<0.005, Fisher PLSD. (G) Relative amounts of activities from 1 h to 25 h after hypoxia corresponding to (B)-(E). Interaction between DHx effect and MP effect was evaluated by two-way ANOVA (vehicle-normoxia; n=9, MP-normoxia; n=9, vehicle-hypoxia; n=11, MP-hypoxia; n=11; ***p<0.0001). †p<0.001, ††p<0.0005, Fisher PLSD. (H, I) Relative variations of daily activities. (H) corresponds to (B, C). (I) corresponds to (D, E). Open arrows indicate vehicle or MP at the beginning of DHx. Closed circles with solid lines and open circles with broken lines indicate relative values of every 2 h activities (mean ± SEM) of MP and vehicle treated mice, respectively. The black and white bars under the values represent the LD cycles. The effects of MP on 24h activity variations of animals exposed to normoxia (H; n=9 vs n=9) and DHx (I; n=11 vs n=11) were evaluated by two-way ANOVA (****p<0.0001). ⍰⍰⍰⍰p<0.0001, ⍰⍰p<0.005, ⍰p<0.01, ⍰⍰⍰⍰p<0.0001, Fisher PLSD.

### DHx reduced adenosine (Ado) in the forebrain and morning inhibition of Ado changed behavior similar to DHx

Microarray analysis revealed DHx increased Ecto-5’-nucleotidase (Nt5e) and adenosine deaminase (Ada) mRNAs in the forebrain (Figure 4A). Nt5e converts adenosine monophosphate into adenosine (Ado), a physiological sleep factor [36]. Ada converts Ado into inosine (Ino). qPCR analysis showed significant Ada mRNA increase (Figure 4B). HPLC analysis clarified Ado and Ino reductions by DHx (Figure 4C). To know the role of Ado reduction, we administered caffeine (15 mg/kg), adenosine A1 and A2A receptor (A1AR, A2AAR) antagonist, to WT at ZT0 (morning caffeine) (Figure 4D, E). After acute activity increase (ZT2-4) (Figure 4F), following 24 h (ZT4-4) activity amount reduced (Figure 4G). Interestingly, morning caffeine reduced the AP activity (ZT12-18, ZT22-0) and increased the AEP activity (ZT0-4) (two-way ANOVA) similar to DHx (Figure 4F). Caffeine (15 mg/kg) administrated at ZT11.5 (evening caffeine) (Supplementary Figure S3A, B) acutely increased activity (ZT12-14) (Supplementary Figure S3C) but did not change following 24 h (ZT14-14) activity amount (Supplementary Figure S3D) and variation (two-way ANOVA) (Supplementary Figure S3C). Next, we administrated caffeine (30 mg/kg) or vehicle to CryDKO 6 h after transition from LD to DD (Figure 4H, I) because HDHT induced wavelike response. After acute increase by caffeine, mean activity value decreased to 18-20 h time point in DD, increased for next 6 h to 26-28 h time point, again decreased to 32-34 h time point (one-way ANOVA) (Figure 4K). Vehicle had no effect in this period (Figure 4J) (one-way ANOVA). We next administrated A2AAR antagonist, istradefylline (3.3 mg/kg), or vehicle to CryDKO at the transition from LD to DD (Figure 4L, M) because A2AAR mediate the arousal effect of caffeine [37]. Istradefylline acutely increased activity amount (2-4 h after treatment) (Figure 4O). After acute increase by istradefylline, activity value decreased to 6-8 h time point in DD, increased to 14-16 h time point, again decreased to 26-28 h time point (one-way ANOVA) (Figure 4O). Vehicle had no effect in this period (Figure 4N) (one-way ANOVA). Results indicated Ado reduction by DHx contributed to DUR and wavelike response of activity by hypoxia.

**Figure 4.**
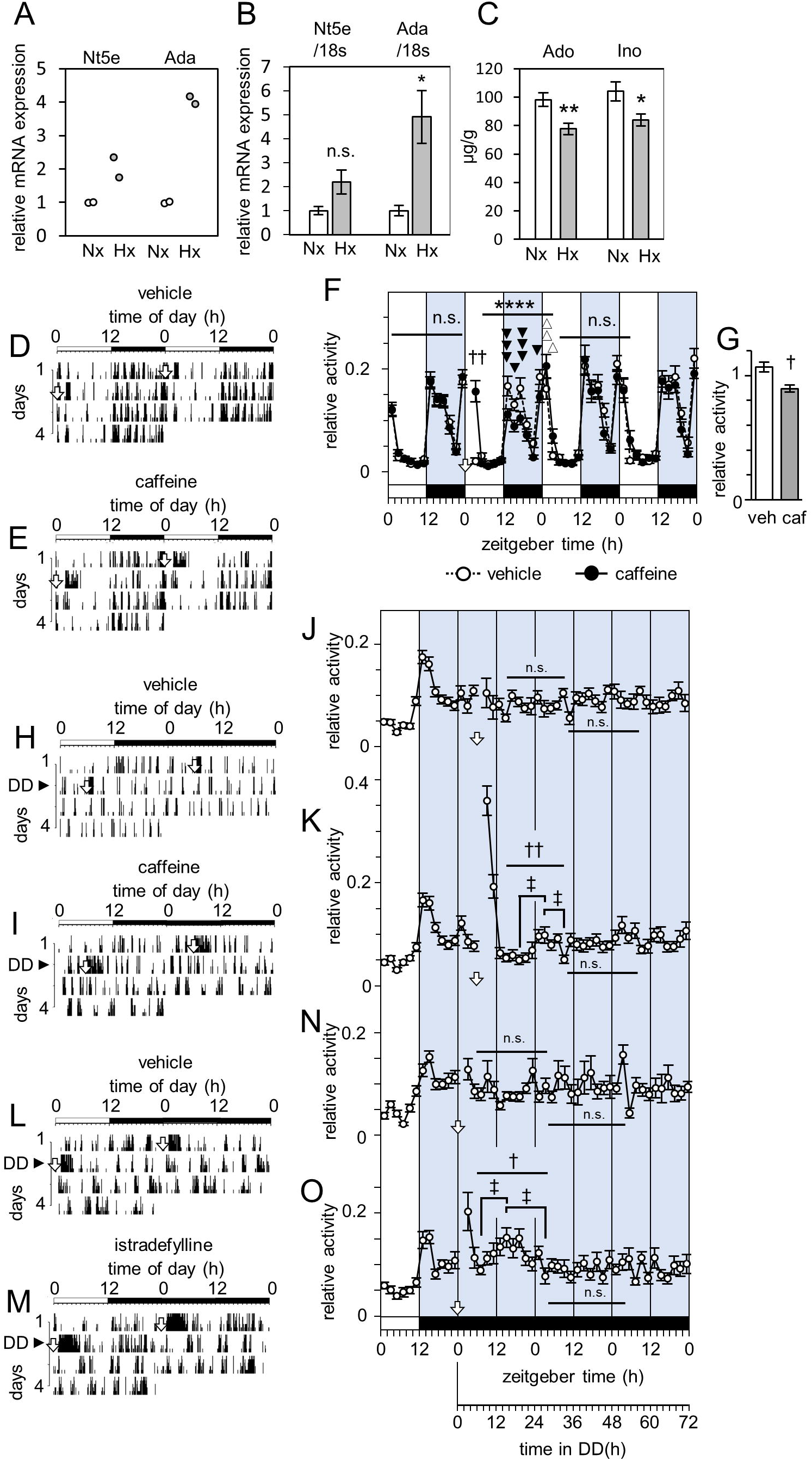
Involvement of adenosine on activity change by DHx. (A-C) *Nt5e* and *Ada* mRNA expressions and Ado and Ino amounts in mice forebrains at the end of DHx (Hx; ZT9) or normoxic control (Nx; ZT9). (A) Quantified values of microarray (ICR; n=2). (B) qPCR values, normalized by 18s expression values (ICR; n=4). (C) Ado and Ino amounts per brain tissue (μg/g) (WT; n=6). *p<0.05, **p<0.01, Student’s t-test. (D-O) Effect of adenosine inhibition on the following activity amount and variation of WT and CryDKO. (D, E) Representative double plotted actograms of WT under LD. The black and white bars above the recordings represent the LD cycles. Open arrows indicate vehicle (D) or morning caffeine (15 mg/kg) (E) at ZT0. (H, I, L, M) Representative double plotted actograms of CryDKO under transition from LD to DD. The black and white bars above the recordings represent the LD and DD. Black triangles indicate the day of moving from LD to DD. Open arrows indicate vehicle (H) or caffeine (30 mg/kg) (I) after 6h from the beginning of DD or vehicle (L) or istradefylline (3.3 mg/kg) (M) at the beginning of DD. (F) Relative amounts of daily activity variations which correspond to (D, E). Open arrow indicates administrations of caffeine or vehicle at ZT6. Closed circles with solid lines and open circles with broken lines indicate relative values of every 2 h activities (mean ± SEM) of caffeine-(n=17) and vehicle-(n=17) treated mice respectively. The black and white bars under the values represent the LD cycles. The acute (2-4h after treatment) effect of caffeine (F) was evaluated by Student’s t-test (†† P<0.0001). Following 24h (ZT4-4) effects on behavior variations were evaluated by two-way ANOVA (**** P<0.0001). ⍰⍰⍰p<0.001, ⍰p<0.05, ⍰⍰p<0.01, ⍰p<0.05, Fisher PLSD. (G) 24 h activity amounts of caffeine (caf, n=17) and vehicle (veh, n=17) treated WT after acute increase (ZT4-4) was evaluated by Student’s t-test. (†P<0.005). (J, K, N, O) Relative amounts of daily activity variations which correspond to (H, I, L, M). Open circles with solid lines indicate relative values of every 2 h activities (mean ± SEM) of caffeine-(n=35) (K) or vehicle-(n=35) (J) treated mice or istradefylline-(n=32) (O) or vehicle-(n=32) (N) treated mice. After direct activity increase by caffeine or istradefylline, activity up-and-downs (caffeine; 14-34 h in DD, istradefylline; 6-28 h in DD) were evaluated by one-way ANOVA (†† p<0.005, † p<0.05).‡ p<0.005, Fisher PLSD. Open arrows indicate drug or vehicle administrations. The black and white bars under the values represent the LD and DD.

### Chronic caffeine treatment induced circasemidian (~12h) and circadian activity rhythms in molecular clock deficient mice

If this wavelike response by single Ado inhibition is a damped oscillation, chronic caffeine treatment may induce continuous oscillation, activity rhythm. Previously, we and other groups showed chronic methamphetamine (MAP) treatment induced circadian activity rhythm which was SCN and molecular clock independent [29, 38–44]. MAP activate dopaminergic neurotransmission as dopamine reuptake inhibitor [45]. D1 and D2 dopamine receptors (D1DR and D2DR) form heteromers with A1AR and A2AAR in the striatum. Ado antagonizes to actions of dopamine at A1AR/D1DR and A2AAR/D2DR heteromers [46, 47]. Because these findings suggest Ado blocker acts similarly with MAP, we started caffeine administration in drinking water (0.5g/L) to CryDKO under LD (Figure 5A). 9 days later, mice were moved to DD. In 16 animals (Supplementary Table S2), 15 animals expressed circasemidian (8.75-17.16 h period) activity rhythms (Figure 5B, D, S4A, B) and 13 animals expressed rhythms with around and longer than 24h periods (20.75-39.41 h period; Figure 5C, S5A, B). One animal showed neither circasemidian nor circadian rhythm for 37 days. Circasemidian and circadian rhythms free ran with changing periods (Supplementary Table S2). Amount of spontaneous caffeine intake was 2.8±0.19 mg/day/animal. After caffeine withdrawal, all CryDKO became arrhythmic. Circasemidian and circadian rhythms by chronic caffeine suggested damped oscillation by single Ado reduction or inhibition drives wave like response and DUR.

**Figure 5.**
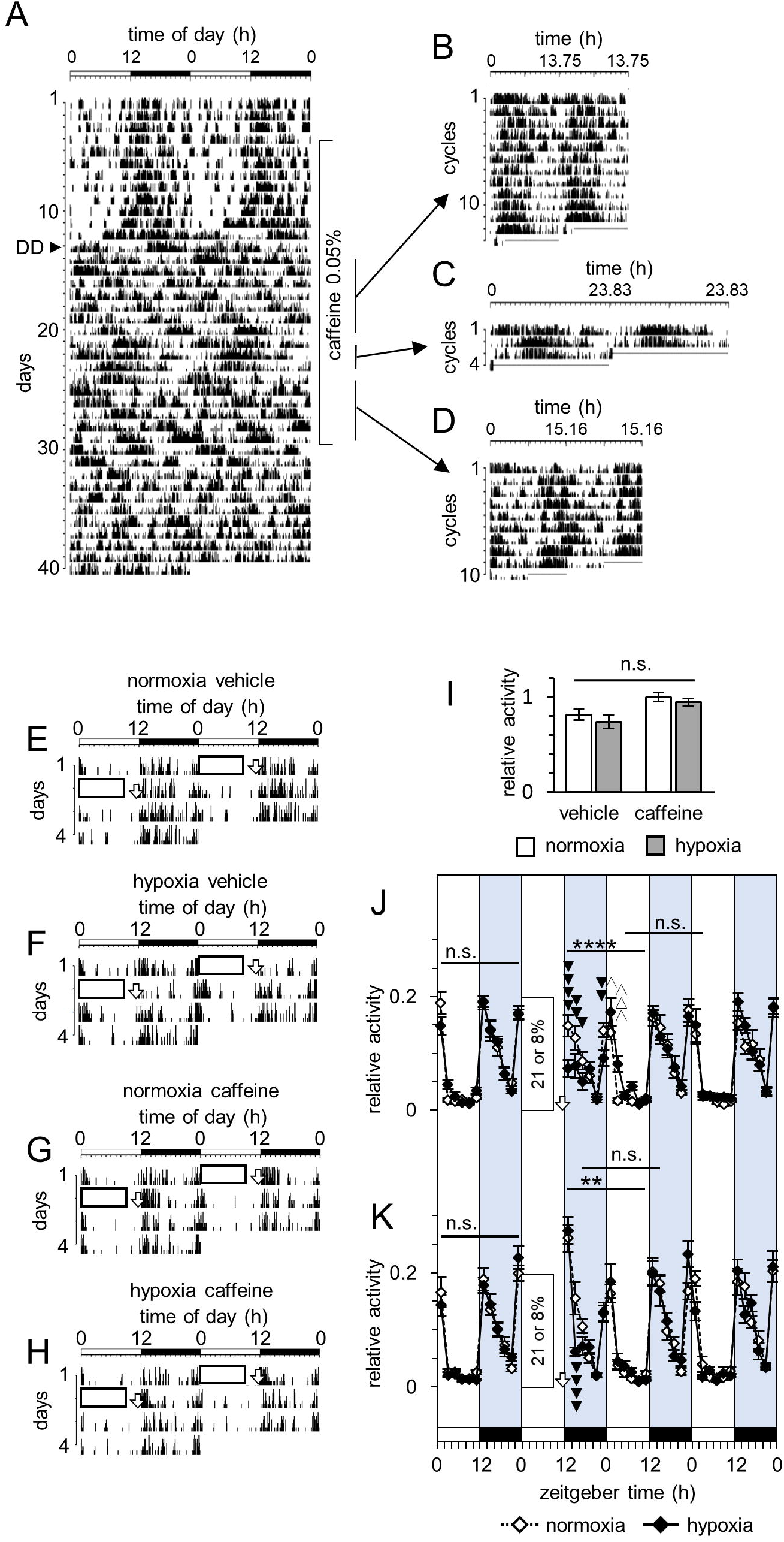
Circasemidian and circadian rhythms of CryDKO by chronic caffeine treatment and effects of evening caffeine on DHx induced WT activity changes. (A-H) Representative double plotted actograms of CryDKO (A-D) and WT (E-H). (A) and (E-H) are plotted on a 2 × 24-h time base. (B), (C) and (D) are plotted on 2 × 13.75-h (7d), 2 × 23.83-h (3d) and 2 × 15.16-h (6d) time bases respectively. The black and white bars above the recordings represent the LD cycles. Black triangle indicate the day of moving from LD to DD (A). Rectangles in the actograms show normoxic control (21%) (E, G) or DHx (8%) (F, H). Open arrows indicate vehicle (E, F) or caffeine (15 mg/kg) (G, H) administrations at ZT11.5. (I) Relative amount of 24 h activities from 0.5 h after caffeine or vehicle administration (ZT12-12). Interaction between DHx effect and caffeine effect was evaluated by two-way ANOVA (normoxia-vehicle; n=12, hypoxia-vehicle; n=12, normoxia-caffeine; n=11, hypoxia-caffeine; n=11). (J, K) Relative amounts of activity variation. (J) and (K) correspond to (E, F) and (G, H) respectively. Open arrows indicate administrations of vehicle (J) or caffeine (15 mg/kg) (K) at ZT11.5. Closed diamonds with solid lines and open diamonds with broken lines indicate relative values of every 2 h activities (mean ± SEM) of DHx-exposed and normoxic control mice respectively (G; n=12 and n=12, H; n=11 and n=11). The black and white bars under the values represent the LD cycles. The effects of DHx on activity variations of vehicle (J) or caffeine (K) treated animals were evaluated by two-way ANOVA (** P<0.005, **** P<0.0001). ⍰⍰⍰⍰p<0.0001, ⍰⍰p<0.01, ⍰p<0.05, ⍰⍰⍰p<0.0005, ⍰p<0.05, Fishers PLSD

### Evening caffeine attenuated the behavior change by DHx

If damped oscillation by Ado inhibition drives wavelike response and DUR, it is possible that Ado inhibition at different timing interferes and attenuates DUR by DHx. Next, we administered evening caffeine or vehicle (ZT11.5) after DHx (Figure 5E-H). There was no interaction between DHx and caffeine effect on 24 h activity amounts from ZT12 (two-way ANOVA) (Figure 5I). DHx changed 24 h activity variations from ZT12 in evening vehicle treated mice (two-way ANOVA) (Figure 5J). AP activity was lower (ZT12-18, ZT22-0) and AEP activity was higher (ZT0-4) in DHx exposed mice, which reflected DUR by DHx. DHx also changed 24 h variations in evening caffeine treated mice (two-way ANOVA) (Figure 5K). By caffeine, activities of DHx exposed and control mice immediately increased to similar level (ZT12-14). Soon later, activity of DHx exposed mice became lower (ZT14-16). Interestingly, following 24 h (ZT16-16) activity variations were not different between these two groups (two-way ANOVA) (Figure 5K). These results showed that evening caffeine attenuated DUR by DHx.

## Discussion

We observed behavioral responses which are comparable to symptoms of COMISA in DHx exposed mice. DHx, AP activity decrease and AEP activity increase in mice corresponded to nighttime sleep apnea, daytime sleepiness and co-occurring nighttime insomnia in COMISA respectively. Especially, AEP activity increase corresponded to difficulty in falling asleep at sleep onset in COMISA [13, 48]. Different from middle insomnia, initial insomnia cannot be direct result of apnea and our animal model suggests apnea (hypoxia) of last night is important for onset insomnia. In CryDKO, DHx also induced DUR and HDHT induced down and up wavelike activity response in DD. Molecular clock independent wavelike response may drive sleepiness and insomnia in COMISA.

Our animal model shows endogenous GC (corticosterone) response alleviates DHx effect (Figure 3). Hypoxia also increases GC (cortisol) in humans [49, 50]. If increased cortisol acts similarly, GC administration against hypoxia may be reasonable. Indeed, GC is already used for the prevention and treatment of Acute Mountain Sickness [9]. Hyperarousal is considered to mask daytime sleepiness and cause insomnia at night in COMISA [13]. Increase in complaints of daytime excessive sleepiness and decrease in complaints of insomnia in severe OSA support this idea [51]. GC increase by activation of hypothalamic-pituitary-adrenal (HPA) axis is postulated to cause hyperarousal in COMISA although it is difficult to show the evidence [15, 52]. In our animal model, repression of GC increase enhanced AP activity reduction but did not affect AEP activity increase by DHx (Figure 3), which suggests GC (cortisol) increase by hypoxia acts to reduce excessive daytime sleepiness but less affects insomnia in COMISA.

DHx reduced Ado in the forebrain. Morning Ado inhibition by caffeine induced DUR same as DHx. Caffeine and istradefylline induced molecular clock independent wavelike response similarly with HDHT. Results indicate Ado reduction by DHx induce DUR in mice and provide a possibility that Ado reduction by apnea-hypoxia induce daytime sleepiness and nighttime insomnia in COMISA. We also consider Ado inhibition causes responses like COMISA. Single caffeine intake before going to bed causes sleep disturbance [53–55]. But, following daytime rebound sleepiness and performance deficits and nighttime alertness increase are not well known. Amount of caffeine intake is correlated with feeling tired in the morning and having difficulty in sleeping in U. S. adolescents [56]. Evening-types consume more caffeine in the evening [57]. These findings suggest that evening caffeine induces daytime sleepiness and nighttime insomnia.

Circasemidian and circadian activity rhythms appeared by chronic caffeine administration in CryDKO. Because caffeine, Ado inhibitor, acts similarly with dopamine at dopamine/Ado receptor heteromer [46, 47], it is possible MAP and caffeine stimulate common system to drive circasemidian/circadian rhythm. Molecular clock independent cell autonomous cellular circasemidian rhythm was reported recently [58]. Circasemidian activity rhythm may be driven at the cellular level. Circasemidian system is hypothesized as a driver of human daytime sleepiness and nap [59, 60]. Our circasemidian activity model may help understanding afternoon sleepiness and nap. The activity down-and-up or up-and-down duration of wave like response by HDHT was ~12 h (Figure 2H), which suggests hypoxia (HDHT and DHx) stimulates this circasemidian system to drive wave like response and DUR.

In contrast to DUR induction by morning caffeine, evening caffeine attenuated DUR by DHx. This result suggests that morning caffeine may improve daytime sleepiness and nighttime insomnia in COMISA. Dopamine reuptake inhibitors, modafinil, armodafinil and solriamfetol, are clinically applied for daytime excessive sleepiness of OSA [61]. Because caffeine accelerates dopamine action at dopamine/Ado receptor heteromers by acting as inhibitor of Ado, it is possible that caffeine improve daytime excessive sleepiness. OSA (sleep disordered breathing) patients consume more caffeine (caffeinated soda [62], Do-It-Yourself Caffeine Audit [63]). A negative correlation is observed between caffeine intake and cognitive impairment in OSA [64]. These findings suggest caffeine improves OSA symptoms. Another report showed negative correlation between OSA marker (3% oxygen desaturation index) and caffeine intake in overweight men [65], which suggests caffeine improves hypoxia of OSA. Mysliwiec and Brock [66] pointed out several possible causes of differential caffeine effects and the possibility that caffeine, in an appropriate dose, may improve OSA. In our model, caffeine acts like DHx or reduces the effect of DHx timing dependently. We think adenosine inhibition chronotherapy, caffeine in an appropriate timing, may improve OSA/COMISA symptoms.

In this mice study, we found DHx induced DUR which mimics COMISA. DUR was based on clock gene independent wavelike response. GC increase by DHx competed AP activity reduction by DHx. Ado decrease by DHx accelerated DUR and wavelike response which is possibly driven by circasemidian system. Evening Ado inhibition after DHx reduced the effect of DHx. Our animal model may advance understanding of COMISA.

## Supporting information

Supporting information includes five figures and two tables.

## Acknowledgements

We thank Dr. K. Yoshikawa and M.Sc. N. Kodama of the Institute of comprehensive medical research Division of advanced research promotion of Aichi Medical University for technical support for microarray and HPLC. We thank Dr. N. Yamaguchi of Department of Pharmacology, Aichi Medical University for technical support for ELISA.

## Funding

This research was funded by MEXT | Japan Society for the Promotion of Science (JSPS) - 22K08246 [Masubuchi], MEXT | Japan Society for the Promotion of Science (JSPS) - 16K08532 [Masubuchi], Akiyama Life Science Foundation [Masubuchi], Suzuken Memorial

Foundation [Masubuchi], Takeda Medical Research Foundation [Masubuchi], Aichi Cancer Research Foundation [Masubuchi]

## Disclosure Statement

There are no financial or conflicts of interest associated with this work.

## Author contributions

S.M. designed research. S.M., T.Y., K.K., K.I., A.O. and S.K. performed experiments. W.N., K.T., Y.H. and T.T. contributed new reagents/analytic tools. S.M., T.Y., A.O., S.K., K.T., and S.T. analyzed data. S.M. wrote the paper.

## Notes

### Competing Interest Statement

The authors have declared no competing interest.

