## Supplementary figures and images for "A mouse model of insomnia with sleep apnea"

### Supplemental Figure 1

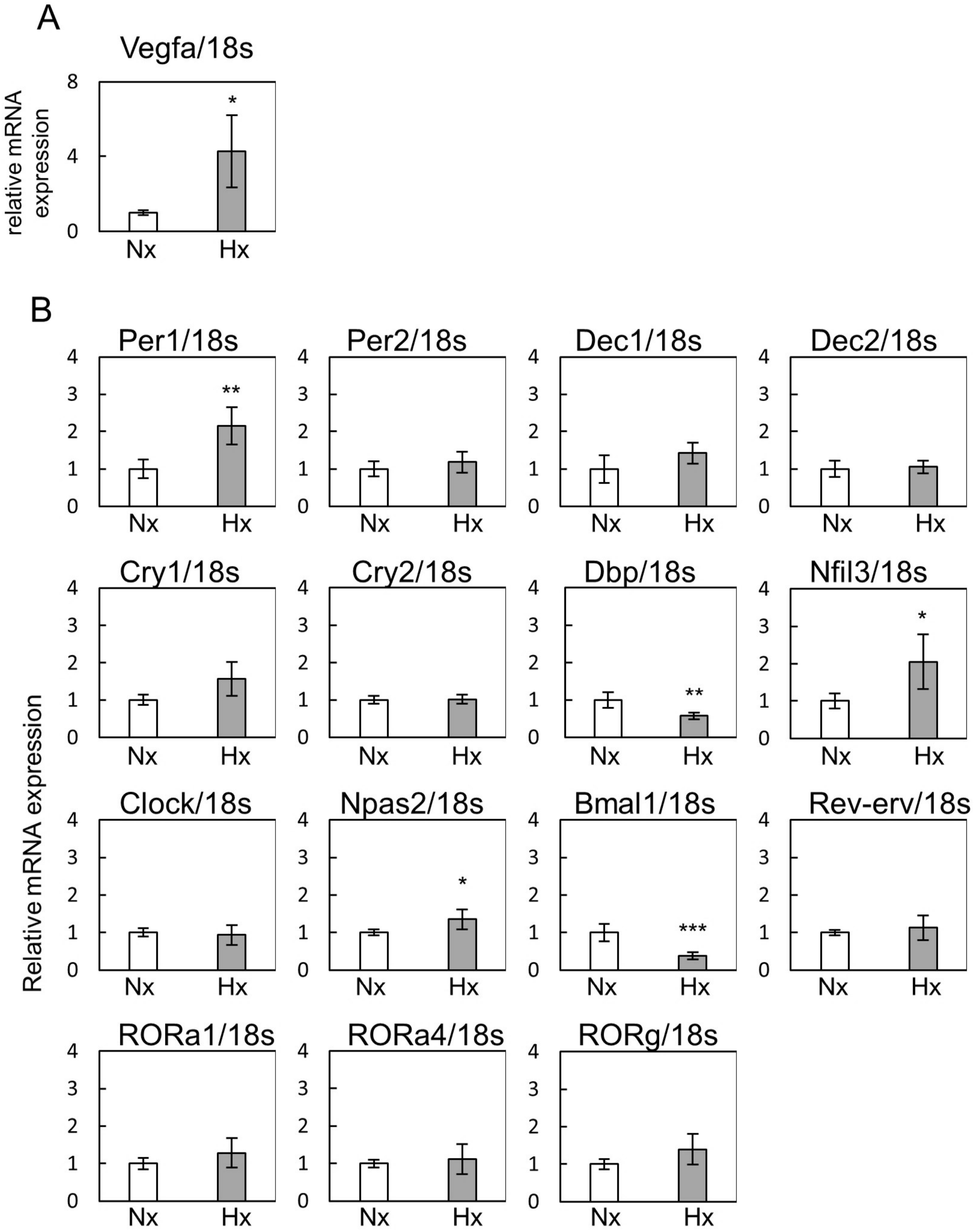


Supplementary Figure S1

### Supplemental Figure 2

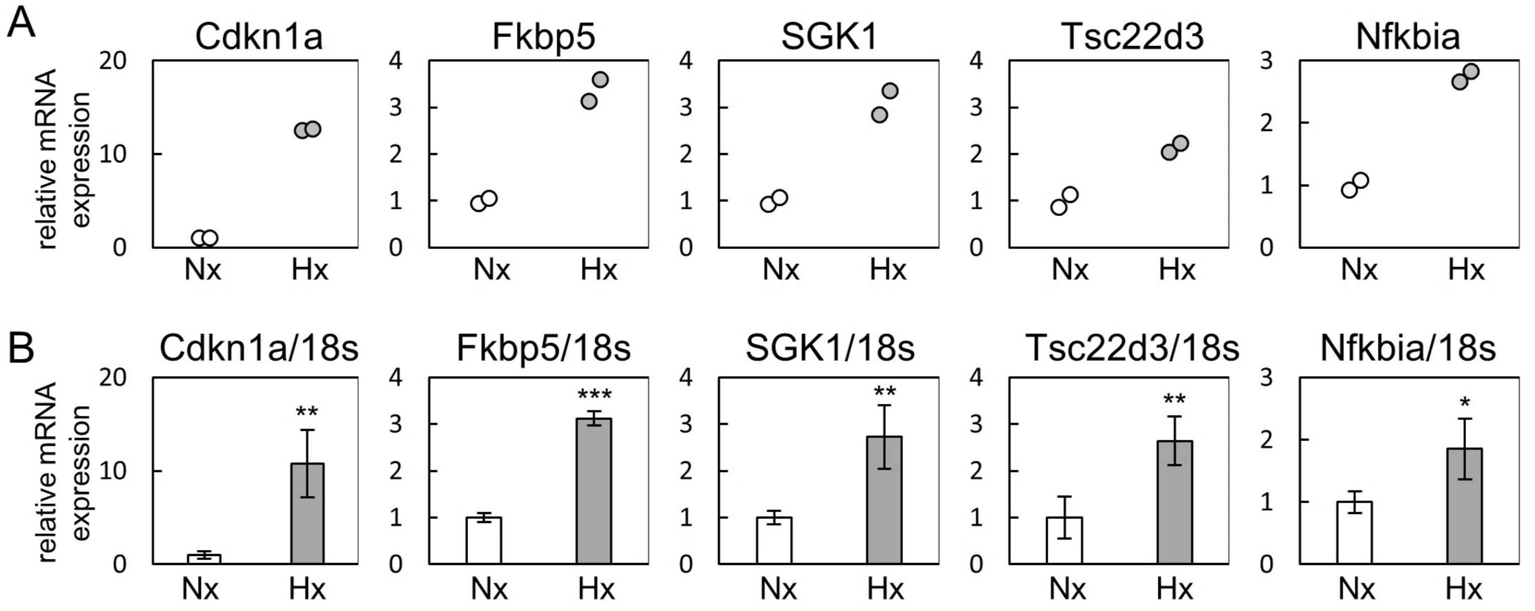


Supplementary Figure S2

### Supplemental Figure 3

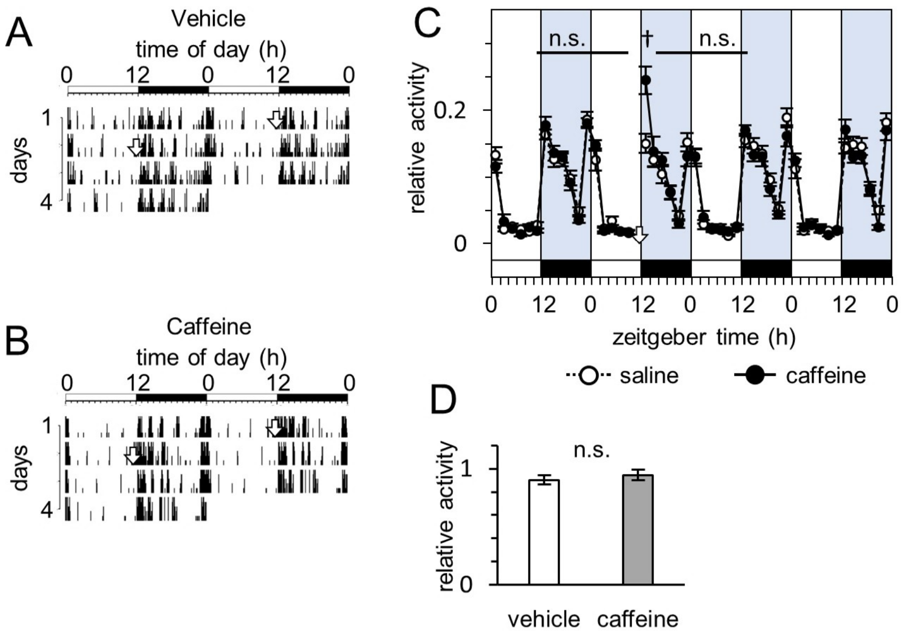


Supplementary Figure S3

### Supplemental Figure 4

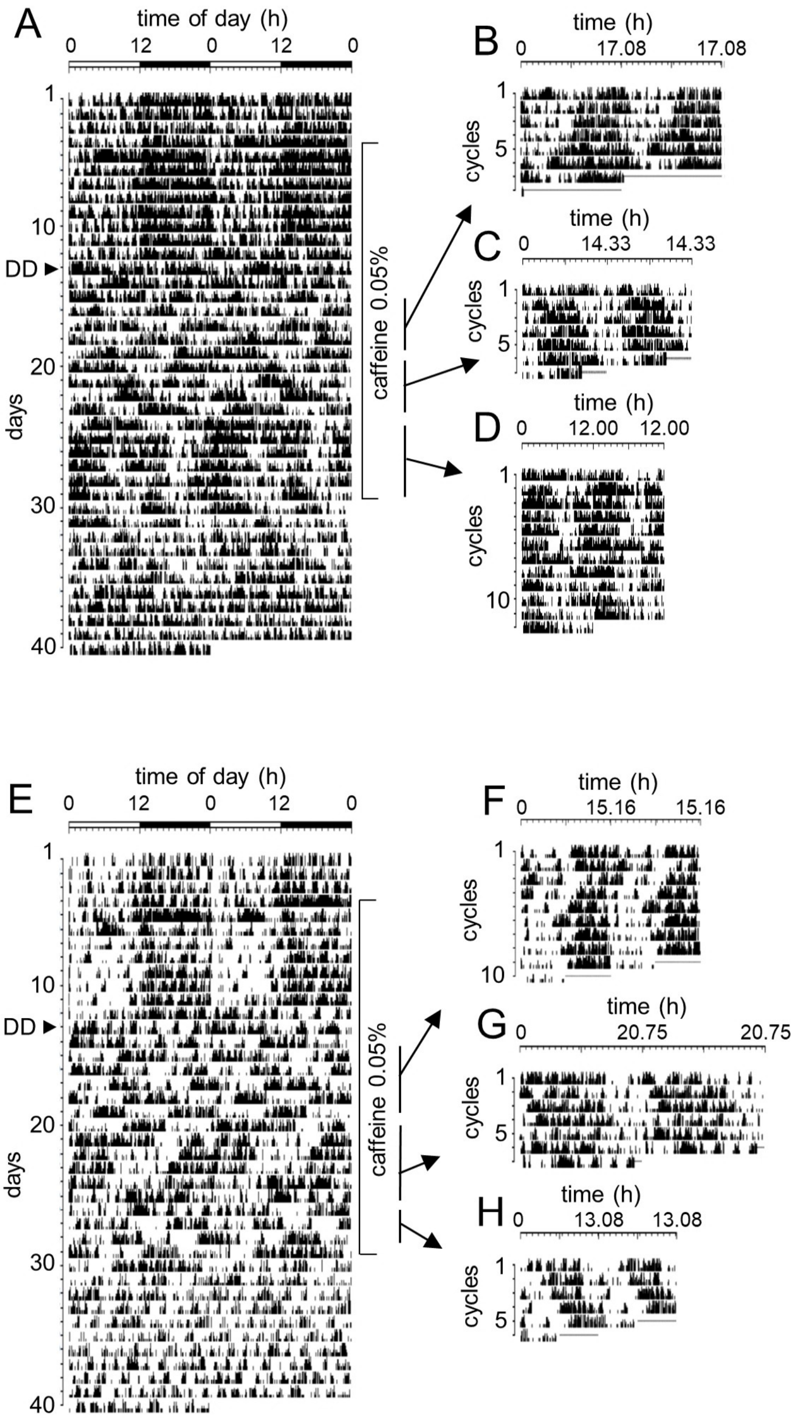


Supplementary Figure S4

### Supplemental Figure 5

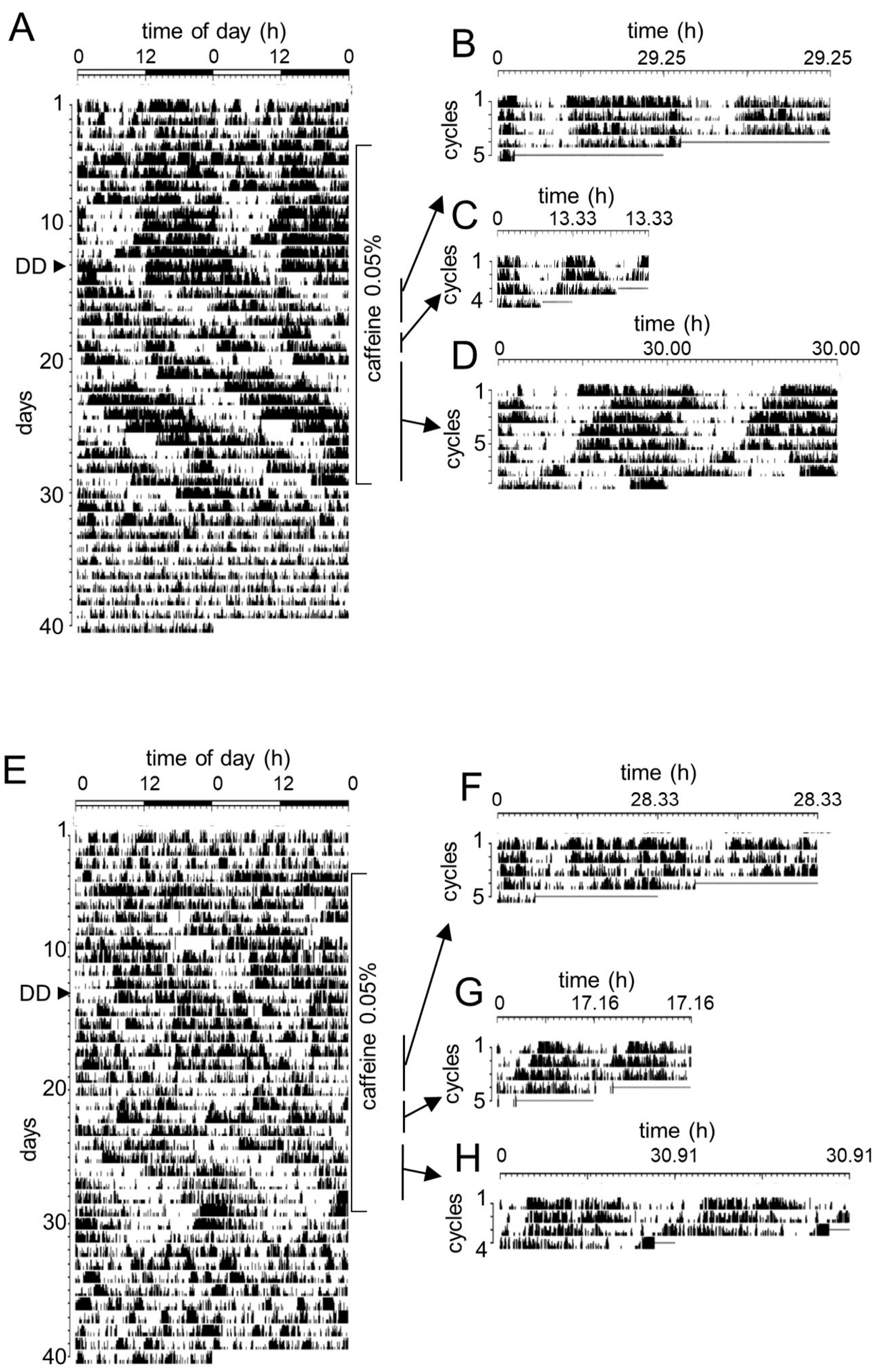


Supplementary Figure S5
