## Supplemental Table 1 for "A mouse model of insomnia with sleep apnea"

**Supplementary Table S1: "Gene expression primer table".**

| ***Gene*** | ***Forward primer*** | ***Reverse primer*** | ***Reference*** |
|---|---|---|---|
| *Vegfa* | TTACTGCTGTACCTCCACC | ACAGGACGGCTTGAAGATG | [S1] |
| *Per1* | CAGGCTAACCAGGAATATTACCAGC | CACAGCCACAGAGAAGGTGTCCTGG | [S2] |
| *Per2* | GGCTTCACCATGCCTGTTGT | GGAGTTATTTCGGAGGCAAGTGT | |
| *Cry1* | CCCAGGCTTTTCAAGGAATGGAACA | TCTCATCATGGTCATCAGACAGAGG | |
| *Cry2* | GGGACTCTGTCTATTGGCATCTG | GTCACTCTAGCCCGCTTGGT | |
| *Dbp* | AATGACCTTTGAACCTGATCCCGCT | GCTCCAGTACTTCTCATCCTTCTGT | |
| *Clock* | CCTATCCTACCTTGGCCACACA | TCCCGTGGAGCAACCTAGAT | |
| *Npas2* | GTATGCACAGAGCCAAGTGATGTT | TGCTCACTGTGCAGAGATGTTG | |
| *Bmal1* | GCAGTGCCACTGACTACCAAGA | TCCTGGACATTGCATTGCAT | |
| *Rev-erbα* | CGTTCGCATCAATCGCAACC | GATGTGGAGTAGGTGAGGTC | |
| *Dec1* | TGGTGATTTGTCGGGAAGAAATC | CATGCTTCGCCAGGTACTGA | Based on [S3] |
| *Dec2* | GGAATCCCTCATTTGCAAGAGA | TTCAAGCTCCTTTTGGGTTTACA | |
| *Nfil3* | ACGGACCAGGGAGCAGAAC | GGACTTCAGCCTCTCATCCATC | [S4] |
| *RORα1* | GAGGTATCTCAGTCACGAAG | AACAGTTCTTCTGACGAGGACAGG | |
| *RORα4* | TGTGATCGCAGCGATGAAAG | AACAGTTCTTCTGACGAGGACAGG | |
| *RORγ* | ACTACGGGGTTATCACCTGTGAG | GTGCAGGAGTAGGCCACATTAC | |
| *Ada* | AAGCATTTGGCATCAAGGTC | CATAGCCACCACGGTCTTCT | [S5] |
| *Nt5e* | CTGACCCAGGAAGACTACCT | GCTGAGGAAGACATGGCTCT | |
| *Cdkn1a* | TCCCGTGGACAGTGAGCAGTTG | CGTCTCCGTGACGAAGTCAAAG | [S6] |
| *Fkbp5* | GAAGCCGGGAAGCCTAAGTT | CGTGTACTTGCCTCCCTTGA | [S7] |
| *Tsc22d3* | CTTCTCACTCTGCGTGCGT | AAAACAAGCTGCCACTTCAGG | |
| *SGK1* | TGAAGGAAGCAGCAGAAG | CGTTGGAAGGAAGAGAAGA | [S8] |
| *Nfkbia* | TCCTGCACTTGGCAATCATC | AGCCAGCTCTCAGAAGTGCC | [S9] |
| *18s rRNA* | AGTCCCTGCCCTTTGTACACA | CGATCCGAGGGCCTCACTA | [S10] |
