## Supplemental Table 2 for "A mouse model of insomnia with sleep apnea"

Supplementary Table S2: “Activity periods of chronic caffeine treated CryDKO”.

| | activity period (days) in DD | illustration |
|---|---|---|
| CryDKO01 | [–] (15d), 11.58(3d), 23.41(4d), 21.83(4d) | |
| CryDKO02 | [–] (4d), 11.91(5d), 16.58(5d), 9.75(3d) | |
| CryDKO03 | [–] (15d), 29.91(5d), 12.91(6d) | |
| CryDKO04 | [–] (17d), 15.58(4d), 39.41(7d) | |
| CryDKO05 | [–] (37d) | |
| CryDKO06 | [–] (21d), 16.66(4d), [–] (6d) | |
| CryDKO07 | [–] (1d), 29.25(5d), 13.33(2d), 30.00(10d) | Suppl. Figure S5A |
| CryDKO08 | [–] (9d), 8.75(6d), 26.66(5d), 26.41(6d) | |
| CryDKO09 | [–] (1d), 17.08(7d), 14.33(4d), 12.00(5d) | Suppl. Figure S4A |
| CryDKO10 | [–] (1d), 28.33(5d), 17.16(3d), 30.91(5d) | Suppl. Figure S5B |
| CryDKO11 | [–] (15d), 21.5(3d), 38.00(11d) | |
| CryDKO12 | [–] (4d), 15.83(4d), 22.25(5d), 14.91(5d), 30.83(3d), 28.16(4d), 12.66(3d) | |
| CryDKO13 | [–] (2d), 13.75(7d), 23.83(3d), 15.16(6d) | Figure 5A |
| CryDKO14 | [–] (2d), 15.16(6d), 20.75(6d), 13.08(3d) | Suppl. Figure S4B |
| CryDKO15 | [–] (14d), 13.16(3d), 26.33(4d) | |
| CryDKO16 | [–] (28d), 13.16(3d), 37.5(6d) | |
