## Supplemental Figure legends for "A mouse model of insomnia with sleep apnea"

**Supplementary Figure S1 Effect of DHx on forebrain gene expressions (related to Figure 1).**

(A, B) *Vegfa* mRNA expressions (A) and Clock genes and clock controlled genes mRNA expressions (B) of ICR mice forebrains at the end of DHx (Hx; ZT9, n=4) and control (Nx; ZT9, n=4) by qPCR. mRNA expression values were normalized with 18s expression values. *$p<0.05$, **$p<0.01$, ***$p<0.005$, Student's t-test.

**Supplementary Figure S2 Effect of DHx on glucocorticoid responsive forebrain gene expressions (related to Figure 3).**

(A, B) Increases of mRNA expressions of glucocorticoid responsive genes in forebrains of ICR mice at the end of DHx (Hx; ZT9) and normoxic control (Nx; ZT9). Quantified values of microarray (ICR; n=2) (A). mRNA expression values by qPCR which are normalized with 18s expression values (ICR; n=4) (B). *$p<0.05$, **$p<0.005$, ***$p<0.0001$, Student's t-test.

**Supplementary Figure S3 Effects of evening caffeine (ZT11.5) on WT activities (related to Figure 4 and Figure 5).**

(A, B) Representative double plotted actograms of vehicle (A) or caffeine (B) treated mice. The black and white bars above the recordings represent the LD cycles. Open arrows indicate or caffeine (15 mg/kg) treatment at ZT11.5. (C) Activity variations correspond to (A) and (B). Open arrows indicate evening caffeine (15 mg/kg) or vehicle treatment at ZT11.5. Closed with solid lines and open circles with broken lines indicate relative values of every 2 h activities (mean ± SEM) of caffeine (15 mg/kg, n=14) and vehicle (n=14) treated mice respectively. The black and white bars under the values represent the LD cycles. The acute (ZT0-2) effect of and effect on following 24h (ZT2-2) behavior variation were evaluated by Student's t-test (†P<0.01) and two-way ANOVA respectively. (D) 24 h activity amounts (ZT2-2) of caffeine (n=14) and vehicle (n=14) treated WT after acute activity increase were evaluated by Student's t-test.

**Supplementary Figure S4 Representative circasemidian rhythms of CryDKO by chronic caffeine treatment (related to Figure 5).**

(A-H) Representative double plotted actograms of CryDKO. (A) and (E) are plotted on a 2 X 24-h time base. (B), (C), (D), (F), (G) and (H) are plotted on 2 X 17.08-h (7d), 2 X 14.33-h (4d), 2 X 23.75-h (5d), 2 X 15.16-h (6d), 2 X 20.75-h (6d) and 2 X 13.08-h (3d) time bases respectively. The black and white bars above the recordings represent the LD cycles. Black triangles indicate the day of moving from LD to DD (A, E).

**Supplementary Figure S5 Representative circadian and circasemidian rhythms of CryDKO by chronic caffeine treatment (related to Figure 5).**

(A-H) Representative double plotted actograms of CryDKO. (A) and (E) are plotted on a 2 X 24-h time base. (B), (C), (D), (F), (G) and (H) are plotted on 2 X 29.25-h (5d), 2 X 13.33-h (2d), 2 X 30.00-h (10d), 2 X 28.33-h (5d), 2 X 17.16-h (3d) and 2 X 30.91-h (5d) time bases respectively. The black and white bars above the recordings represent the LD cycles. Black triangles indicate the day of moving from LD to DD (A, E).
